# The reach of reactivation: Effects of consciously-triggered versus unconsciously-triggered reactivation of associative memory

**DOI:** 10.1101/2023.07.26.546400

**Authors:** Amir Tal, Eitan Schechtman, Bruce Caughran, Ken A Paller, Lila Davachi

## Abstract

Newly formed memories are not passively stored for future retrieval; rather, they are reactivated offline and thereby strengthened and transformed. However, reactivation is not a uniform process: it occurs throughout different states of consciousness, including conscious rehearsal during wakefulness and unconscious processing during both wakefulness and sleep. In this study, we explore the consequences of reactivation during conscious and unconscious awake states. Forty-one participants learned associations consisting of adjective-object-position triads. Objects were clustered into distinct semantic groups (e.g., multiple fruits, vehicles, musical instruments) which allowed us to examine the consequences of reactivation on semantically-related memories. After an extensive learning phase, some triads were reactivated consciously, through cued retrieval, or unconsciously, through subliminal priming. In both conditions, the adjective was used as the cue. Reactivation impacted memory for the most distal association (i.e., the spatial position of associated objects) in a consciousness-dependent and memory-strength-dependent manner. First, conscious reactivation of a triad resulted in a weakening of other semantically related memories, but only those that were initially more accurate (i.e., memories with lower pre-reactivation spatial errors). This is similar to what has been previously demonstrated in studies employing retrieval-induced forgetting designs. Unconscious reactivation, on the other hand, benefited memory selectively for weak cued items. Semantically linked associations were not impaired, but rather integrated with the reactivated memory. Taken together, our results demonstrate that conscious and unconscious reactivation of memories during wakefulness have qualitatively different consequences on memory for distal associations. Effects are memory-strength-dependent, as has been shown for reactivation during sleep. Results support a consciousness-dependent inhibition account, according to which unconscious reactivation involves less inhibitory dynamics than conscious reactivation, thus allowing more liberal spread of activation. Our findings set the stage for additional exploration into the role of consciousness in memory structuring.

## Introduction

Most of the events we experience will be forgotten, but some will transform into lasting memories ^1^. Whether or not the memory for a recent experience is preserved relates both to the progression of forgetting and to an extended process that transpires after the experience of an event has ended, namely consolidation. *Consolidation* refers to post-encoding stabilization and reorganization of memories and, for episodic memory is thought to involve hippocampal-cortical brain networks ^2^. A critical mechanism supporting the process of consolidation may be the post-encoding replay or reactivation of memories ^3^. Although little is known about the relationship between consciousness and memory reactivation, it is assumed that we can be either unaware or aware of a transpiring reactivation event and that it can occur during either sleep or wakefulness ^3,4^. The unique mechanisms that play out in these different circumstances are presently unclear ^5^.

Sleep possesses some unique characteristics that may render it uniquely suited for optimal reactivation. Sleep is relatively sheltered from external stimuli and involves an oscillatory milieu that may optimize communications between the cortex and subcortical regions ^1,4,6,7^. However, another fundamental characteristic of memory reactivation during sleep, that has received little attention, is that it apparently occurs outside the realm of conscious awareness. During wakefulness, spontaneous reactivations driving memory consolidation occur during offline periods, characterized by a lack of a task or goal and by reduced outward attention ^5,8–11^. Recent work has found that briefly cueing memories while participants were performing an unrelated task benefited later memory to a greater extent for participants who were less vigilant and engaged (as reflected by longer response times on the unrelated task) and less aware of the cueing, suggesting that the benefits of memory reactivation could be greater during offline as compared to online states for certain kinds of memories ^12^. These findings, taken together, raise the question of how conscious access to reactivated content moderates the consequences of memory reactivation.

It is important to consider the mechanistic distinctions between conscious and unconscious reactivation. Conscious memory retrieval is often characterized as a competitive process in which one memory is selected and brought into conscious awareness while competing alternatives are suppressed and made less accessible ^13–16^. This process of selection improves directed access to the desired memory ^17^ and may even be useful for reducing interference during future similar retrieval events ^18,19^. This form of directed memory retrieval is thought to be driven by prefrontal-controlled inhibition ^20,21^. Inhibition of non-targets may rely, therefore, on the conscious activation of the target memory. In reactivation scenarios in which no memory is selected for conscious retrieval, competing activations are spared suppression and allowed to persist. While this may be harmful to goal-directed behavior, it may benefit memory restructuring in the brain by permitting more distal associations to be strengthened. Accordingly, we hypothesized that unconsciously-triggered reactivations will therefore lead to greater benefits to more distal and related associations, and may also lead to increased binding between them.

To address this question, in this study we contrasted the effects of conscious versus unconscious reactivation on later memory for triads. Our design allowed us to investigate the effects these reactivations had on associations that were either proximal or distal to the reactivated memory (vertical associative spread), and also on memories that were semantically related to the reactivated memory (horizontal associative spread). To do so, we identified perceptual thresholds in each participant to reactivate compound memories, made of a adjective-object-position triad, by presenting the adjectives below or above the threshold for consciously experiencing them. Importantly, we controlled reactivation using cues to selectively reactivate only a subset of recent memories. This procedure builds on targeted memory reactivation, a technique used to bias reactivation during sleep via unobtrusive presentation of learning-related stimuli ^22,23^, previously adapted for awake reactivation as well ^12,24^. We then tested memory for proximal adjective-object associations and for more distal object-position associations, both in reactivated memories and in semantically linked memories that were not themselves cued.

We hypothesized that unconscious reactivation while awake might resemble sleep reactivation and thus differ from conscious reactivation in two major ways: (1) it will elicit more liberal associative spread, not imposing suppression on distal and related semantic memories; and (2) it will be particularly beneficial for weak memories ^10,12,25–28^. On the other hand, we hypothesized that conscious reactivation may have inhibitory consequences on related memories, replicating retrieval-induced forgetting (RIF; Anderson, 2003). Lastly, we hypothesized that unconscious reactivation may also lead to increased integration with related memories, due to their more liberal spread and, hence, concurrent reactivation ^29^. Our results support these hypotheses, suggesting that activation and inhibition dynamics following reactivation are memory-strength-dependent and consciousness-dependent. While conscious reactivation had an effect on stronger memories – boosting recall of cued associations and impairing memory for related associations – unconscious reactivation benefited weaker memories and facilitated their integration. This diverging pattern of effects suggests that conscious and unconscious reactivations are qualitatively different and may have different unique roles in memory.

## Methods

### Participants

Forty-eight Northwestern University undergraduate students participated in the study for course credit. Data for seven participants were excluded from analysis: three participants did not complete all the critical experimental phases, one participant was excluded due to technical errors, and three participants were excluded due to poor learning of the experimental material (see *Trial and Participant Exclusion*). The final sample included 41 participants (25 identified as female, 16 as male; 38 right-handed) between the ages of 18 and 23 years (M = 18.85, SD = 0.99). The Northwestern University Institutional Review Board approved the procedure, and the experiment was performed in accordance with relevant guidelines and regulations.

### Stimuli

Stimuli in the experiment were used to form three-way associations (“associative triads”); each triad included an adjective, an image of an object, and a spatial on-screen position (Figure 1). Stimuli included 76 adjectives, each of which had 4-6 letters, 1-2 syllables, and were taken from the Medical Research Council psycholinguistics database (https://websites.psychology.uwa.edu.au/school/mrcdatabase/uwa_mrc.htm), with a 500-700 familiarity rating (plus one additional adjective, *scared*). For each participant, 54 adjectives were randomly assigned to triads, an additional 18 were used as novel words in the Reactivation phase, and four were used in practice trials (including one, *short*, that appeared on instruction screens and was always assigned to practice trials).

**Figure 1.**
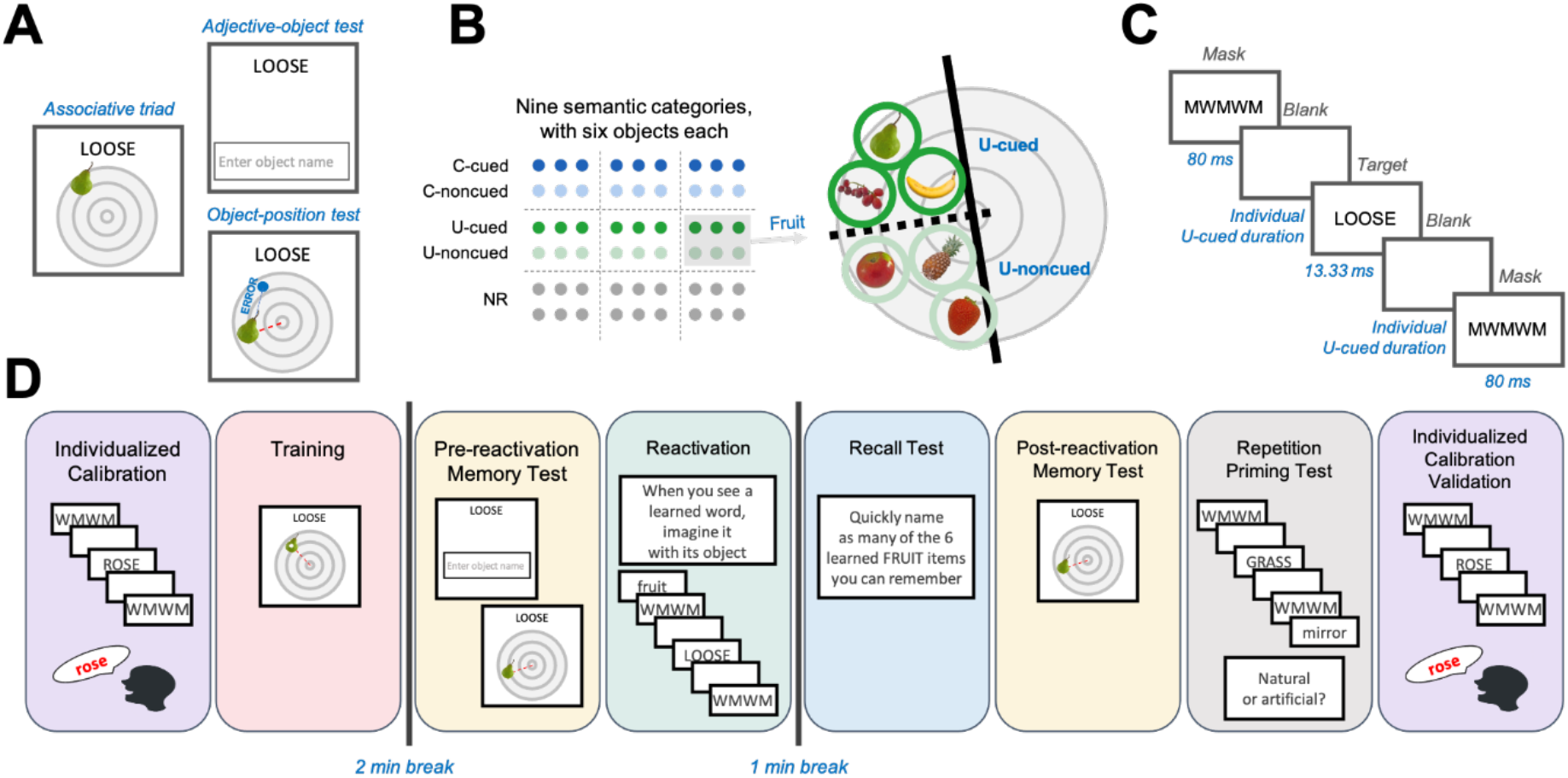
Study design. **A)** Participants learned associative triads comprised of an adjective (e.g., “loose”), an object (e.g., a pear), and a position on a grid. Memory for triads was tested with the adjective-object and object-position tests before and after memory reactivation. In the latter test, recall error is taken as the distance between the object position selected and the veridical position where the object had appeared during training. **B)** Objects in each triad belonged to one of nine semantic categories (e.g., fruit). There were six objects in each category, such that there were 54 triads in total. Half of the memories of six categories (3 × 6 = 18 triads) were reactivated: triads from three categories with conscious reactivation (C-cued) and triads from three categories with unconscious reactivation (U-cued). The three remaining categories were not reactivated (NR). The six objects in each category had systematic spatial positions; the three cued objects were within a 90-degree segment of the grid (dark green circles) and the three noncued objects were within an adjacent 90-degree segment of the grid (light green circles). **C)** In each reactivation trial, an uppercase target adjective (“LOOSE”) was sandwiched between masks (“MWMWMWM”). Between-image blank durations were adjusted to either be brief for subliminal cues or long for supraliminal cues. **D)** Experimental timeline. Participants first underwent a calibration session to determine their individualized perceptual threshold. Then, they were trained to criterion on associative triads. Following a break, memory for adjective-object associations and the object positions was tested. Some of the associations were then reactivated using adjectives as cues (as in panel C). After a break, participants took a recall test for the objects in each category and a test for their spatial positions. Lastly, individualized perceptual thresholds were used in two manipulation checks. Response times were measured for repeated vs non-repeated words, followed by calibration verification as used initially.

Object images were collected from online sources and belonged to nine distinct semantic categories: animals, clothing, fruit, furniture, hospital, games, sports, tools, and vehicles. The ‘games’ category was replaced by the music category for the final 25 participants, as an interim analysis of category-cued recall showed that games and sports categories overlapped and caused confusion. Each of the nine categories included six distinct objects (e.g., apple, banana, grapes, pear, pineapple, strawberry), totaling 54 critical objects. Four additional objects from the plant category were used for practice, one of which was also used for instructions.

For each participant, the 54 objects were each randomly paired with an adjective and were each assigned an on-screen position. These positions were pseudo-random, partly determined by category and condition. Objects from the same category were assigned positions that were confined to one half of the grid (Figure 1B). As nine categories were used, category regions were offset by 40° one from the other and overlapped (each covered 180°). Semi-circles of U and C categories were further divided into two quadrants. The three triads from these categories that were cued during the Reactivation phase (i.e., U-cued and C-cued) were assigned positions in one quadrant, while the three noncued triads (i.e., U-noncued and C-noncued) were assigned positions in the other quadrant (Figure 1B).

For the repetition priming test, 120 nouns were used, each comprised of 4-6 letters; 60 denoted natural objects (e.g., *grass*) and 60 denoted artificial objects (e.g., *mirror*). An additional 112 nouns were used for the Individualized Calibration and Individualized Calibration Validation phases, described below.

Stimuli were presented on a 20.75 × 11.67 inch Dell P2417H screen, controlled by Matlab2020b code using the Psychtoolbox-3 toolbox (Brainard, 1997). The experiment alternated between two presentation settings: 1920 × 1080 pixels at 60 Hz (Slow setting) and 1280 × 1024 pixels at 75 Hz (Fast setting). The Fast setting was used in phases that required masking: Individualized Calibration, Reactivation, Repetition Priming Test, and Individualized Calibration Validation. These phases contained word stimuli only (see the following *Masked Cueing* section for a detailed description of presentation settings during these phases). All other parts of the experiment were done under the Slow setting. For the spatial positioning task, objects were presented such that their long axis extended 150 pixels, overlaid on top of a 700 × 700 pixel image of a circular grid. The entire stimuli set, experimental code and data, can be found online at https://osf.io/fdr8a/.

### Masked Cueing

Our procedure involved multiple phases which included target words presented in between two masks (Figure 1C). We collectively refer to the trials in these phases as masked trials. Depending on the timing of stimuli display, the target word in these trials was sometimes presented subliminally and sometimes supraliminally. These trials make up the Reactivation phase, the core manipulation of the study, but are also used in the Individualized Calibration, Repetition Priming Test and Individualized Calibration Validation phases (Figure 1D).

In masked trials, target words (cues) were presented in uppercase letters between two identical strings of uppercase letters acting as masks: “MWMWMWM”. Each cueing sequence contained, in order: a forward mask, a blank screen, the target word, another blank screen, and a backward mask (i.e., MASK-BLANK-TARGET-BLANK-MASK). Masks were presented for 80 ms and target words for 13.33 ms. The critical manipulation pertained to the duration of the blank screens. When the blank screens had a very short duration, participants could not consciously perceive the target words ^30^. The minimal blank screen duration used was 13.333 ms. To manipulate conscious and unconscious perception, we identified the maximal blank screen duration for each participant that persistently did not produce conscious awareness of the target word (see Procedure). Each individual’s perceptual threshold was then used in some trials to render the target word imperceivable, hence producing unconscious reactivation. In other trials, a longer duration was used to render the target word visible, hence producing conscious reactivation.

### Procedure

The experiment was made of multiple consecutive phases (Figure 1D).

#### 1) Individualized Calibration

The goal of the Individualized Calibration phase was to find the longest blank duration that still renders the target word unreadable (i.e., the perceptual threshold). Specifically, the duration of the blank screens separating target words from surrounding masks in masked trials was manipulated and conscious reports were collected (Fig. 2A). To this end, on each trial participants were shown masked trials which included nouns as target words, and were asked to say out loud the word they saw. If they were unsure or did not consciously perceive any word, they were asked to say the first word that came to mind. An experimenter in the room registered whether each response was correct or incorrect. To identify the perceptual threshold, we used a 1-down-4-up staircase procedure that continuously modified the blank duration: each correct response caused the duration of the blank screen to shorten; four consecutive incorrect responses caused the duration of the blank to lengthen. Both shortening and lengthening were made in steps of 13.33 ms. Blank duration was initialized at 160 ms and was allowed to reach a minimum of 13.33 ms. Calibration ended when the participant’s perceptual threshold was identified, defined as the blank duration that (1) was used in at least 30 trials; and (2) produced less than 10% correct responses.

**Figure 2.**
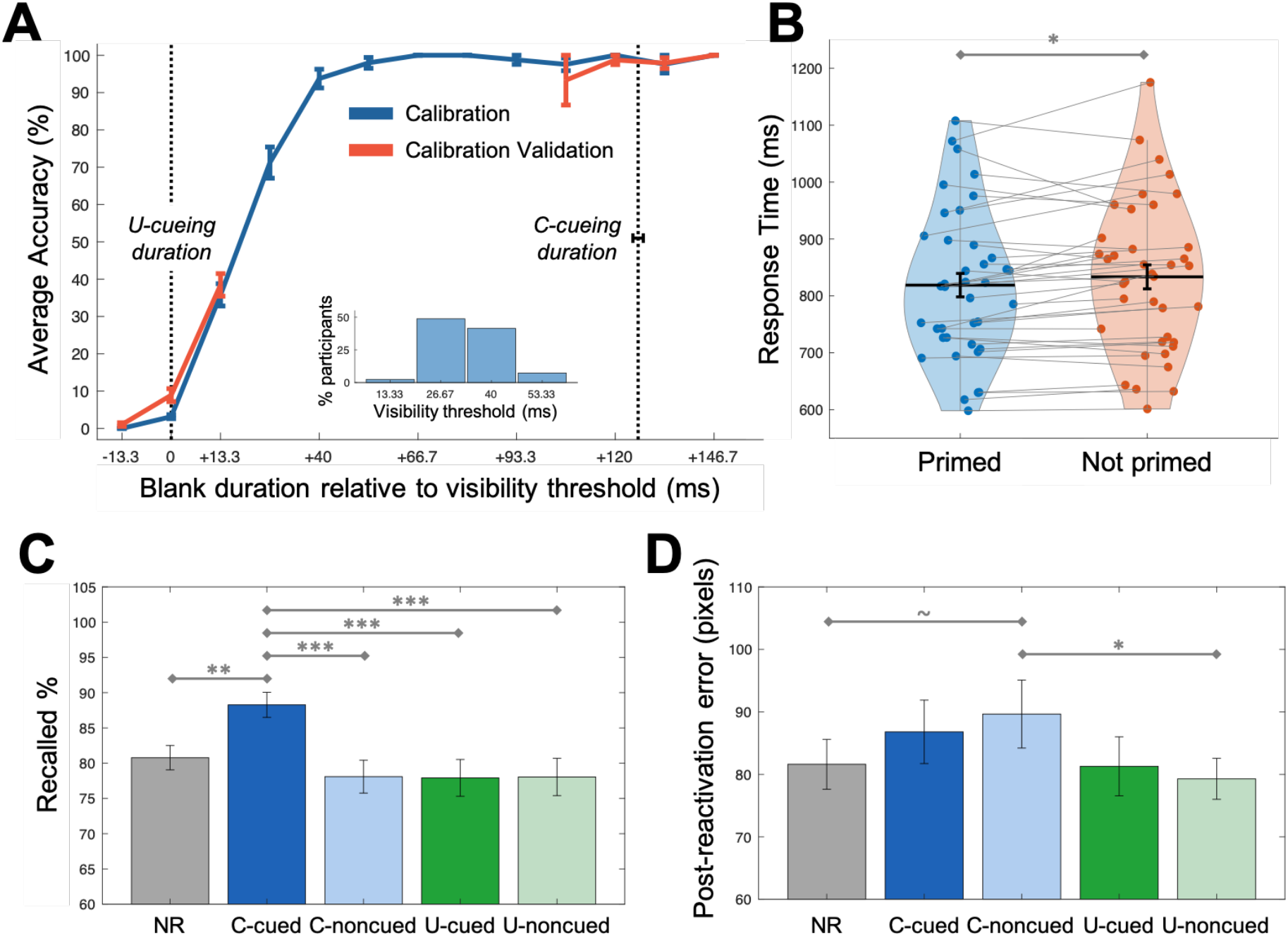
**A)** Individualized visibility calibration. Colored lines indicate the accuracy in naming the masked word as a function of the blank duration relative to the duration parameter chosen for U-cued trials (i.e., the visibility threshold). The blue line reflects the Individualized Calibration session at experiment onset, and the red line reflects the Individualized Calibration Validation session at the end of the experiment. Dotted lines mark the individualized duration that was converged on for U-cued trials (left), and the average gap between it and the 160 ms duration used for C-cued trials (right). Accuracy improved as expected when longer blanks were used, and performance was comparable across sessions. Importantly, word recognition was below 10% when using the U-cued duration, and close to 100% when using the C-cued duration. Inset: distribution of the visibility threshold values across participants **B)** Subliminal priming reduced response times in a semantic judgement task (i.e., repetition priming). This result suggests that the individualized thresholds were sufficient to impact perception. **C)** Category-cued recall was superior for objects from C-cued triads than all other objects. **D)** Post-reactivation position error was larger for C-noncued than for U-noncued and NR objects. In all plots error bars reflect SEM. NR – non-reactivated; C-cued – consciously cued; C-noncued – not cued from groups that contained C-cued items; U-cued – unconsciously cued; U-noncued – not cued from groups that contained U-cued items. ∼ *p* < 0.1, * *p* < 0.05, ** *p* < 0.01, *** *p* < 0.001.

#### 2) Training

The goal of this phase was for participants to learn 54 adjective-object-position triad associations. Objects came from nine distinct semantic categories (see *Stimuli*). Six objects from each category were included. Learning was divided into six blocks. Before the first block, a practice block was administered using an additional category (*plants*) that consisted of four objects only. Each block included training of nine associations. Each pair of consecutive blocks encompassed three categories (e.g., learning block 1 contained three *animal* objects, three *vehicle* objects, and three *tool* objects, and learning block 2 contained the remaining three *animal* objects, three *vehicle* objects, and three *tool* objects). At the beginning of each block, participants were familiarized with each category’s objects before learning their associated adjectives and positions: before each pair of blocks containing three new categories, all items from each of the three categories were displayed on-screen all at once, together with their names, one category at a time. In every three categories that were learned together in the same learning blocks, one category was assigned to each of the three conditions (see Reactivation). This ensured that learning recency was balanced across conditions.

Each learning block began with Guided-training of all associations. In Guided-training, a circular grid appeared at the center of the screen and a fixation cross appeared above it. After 250 ms, the fixation was replaced by an adjective. After 1 s, the object appeared in the center of the grid. After another 500 ms, a white dot appeared on the grid, marking the veridical position for this specific object. Participants were instructed to move the object image (using the mouse) to the position indicated by the white dot and to remember this object-specific position. The Guided-training trial ended when the participant “dropped” the object (by clicking the mouse button) within 20 pixels of the white dot.

After Guided-training of all the associations in the block was completed, Feedback-training began. During this part of the task, the nine adjective-object-position triads were repeated until the learning criteria was met (see below). Feedback-training trials included two consecutive parts. First, the adjective-object association was tested, and then, if a correct response was given, the object-position association was tested (Figure 1A). For adjective-object testing, a fixation cross appeared at the center of the screen for 250 ms, and was then replaced by an adjective (e.g., “SCARED”). Participants were asked to type the name of the object associated with it (e.g., “BANANA”). Participants were allowed to only type in the first four letters of a word to avoid spelling-related errors and save time. A “success” or “failure” tone was then played, along with the correct object image and name which were presented on screen for 1 s. Incorrect trials were then terminated. If correct, an object-position test then commenced: the grid appeared behind the object and participants were asked to drag the object to its correct position and drop it there. Dropping an object within 100 pixels of the veridical position was considered a correct response in this part of the task. Either a green check-mark or a red X appeared in the selected position, and an appropriate “success” or “failure” tone was played. After this feedback, and regardless of whether the response was correct or not, a white dot appeared on the grid indicating the veridical position for the object. This feedback remained on-screen for 1 s before the next trial began.

Testing blocks continued until all associations reached the defined learning criterion. An association was considered learned when both the adjective-object test and the object-position test were marked as correct on two consecutive trials. Once an association was learned, participants no longer had to go through the object-position tests for these triads; Only the adjective-object test was given for the triad from that moment on. Once all nine associations were learned, the block was terminated and the next block began.

#### 3) Pre-reactivation Memory Test

Following a 2-minute break, a memory test of all associations was carried out, to provide a baseline measure of memory prior to reactivation. Testing was similar to testing during the Training phase, with two differences. First, no feedback was given during testing. Second, tests were divided into two parts. Adjective-object associations were tested for all triads, and then object-position associations were tested for all triads. Both parts began with the four practice associations (i.e., linked with the *plant* category), and the remining 54 associations were then tested in random order.

#### 4) Reactivation

After the test, triads were divided to three conditions. Triads belonging to three categories (i.e., 18 adjective-object-position triads) were assigned to the conscious (C) reactivation condition, triads of three categories were assigned to the unconscious (U) reactivation condition, and triads of the remaining three categories were assigned to the non-reactivated (NR) condition. Conditions were assigned such that memory across conditions was matched. This was achieved by dividing the three categories learned in each learning block between the three conditions. This ensured that stimuli of all conditions were encountered at the same times and tested following the same time lag. Then, all options to assign conditions under this constraint were explored, and the configuration producing the minimal variability in pre-reactivation errors across conditions was selected.

Within categories in the reactivated conditions (U and C), only half of the triads (three) were cued during the Reactivation phase. This allowed us to examine later memory for cued and noncued triads from the same category. The cued triads’ spatial positions occupied one quadrant of a category’s semi-circle, whereas the noncued triads’ positions occupied the other quadrant (see *Stimuli*).

In reactivation trials, the triad’s adjective was presented as the target word in a masked trial. For triads in the U-cued condition, the timing constant that was revealed in the calibration phase was used as the duration of the blank screen buffering between the adjective and the surrounding masks, rendering the adjective imperceivable. For triads in the C-cued condition, the duration of the blank screen was always 160 ms, rendering the adjective consciously visible. Triads belonging to the NR condition were not presented in this phase.

During the reactivation phase, participants were instructed to try and identify the target word, presented in uppercase letters. Unlike masked trials presented in the Calibration phase, reactivation trials included an additional screen that preceded the first masked screen. In this screen, a word was presented in lowercase letters for 400 ms (hence clearly visible). It informed participants which category the triad belongs to (e.g., for the adjective “SCARED”, which was paired with a banana, the lowercase word would be “fruit”). Participants were told that the target word may be an adjective that they had learned before, in which case the lowercase word will indicate the category of the object associated with the adjective. If they were able to identify the target word as one of the adjectives, participants should imagine both the adjective and its linked object as vividly as possible (e.g., a scared banana). Participants were told that the main objective of this experiment is to test how imagination affects cognition. Participants were also told that if the target word is either unfamiliar or impossible to see, no action is required. Each masking trial was followed by a 4 s blank screen to provide time for the participants to vividly imagine the associations.

Altogether, 18 adjectives from learned associations were cued during Reactivation: nine U-cued triads and nine C-cued triads (three triads from each of three C/U categories), in addition to 18 novel adjectives used as control which were never seen before in the experiment. Each adjective was reactivated three times. This was done over three blocks, such that each block included a single presentation of all 36 adjectives. Blocks were separated by self-paced breaks. Each block started with buffer trials which consisted of practice triads (i.e., from the *plant* category): the first block started with four such trials (two presented supraliminally and two subliminally); the other blocks started with two supraliminal trials. Altogether, the Reactivation phase comprised 116 trials: 36 adjectives cued 3 times each, and eight practice trials.

#### 5) Object Recall Test

Following a 1-minute break, participants were asked to recall all learned objects. A category name was shown on screen (e.g., “fruit”), and participants were asked to type as many of the objects they remember from that category, in any order. The order in which categories were tested was randomized, except for the practice category (*plants*) which was always tested first. This phase was self-paced and no feedback was provided. Two participants failed to understand the instructions for this phase and their data from this phase was not included in the analysis.

#### 6) Post-reactivation Memory Test

Next, memory for all object-position associations was tested like in the Pre-reactivation Memory Test. Note that unlike the previous phase, adjective-object associations were not tested here.

#### 7) Repetition Priming Test

The goal of this phase was to test whether masking trials presented rapidly (i.e., using an individual’s identified perceptual threshold) produced an unconscious repetition priming effect as has been shown previously (e.g., Dehaene et al, 2001). This phase was added to rule out the possibility that stimuli were presented too rapidly to be processed under U-cued conditions. A repetition priming effect would demonstrate that despite being imperceivable, subliminally cued words were processed.

Participants began each trial by clicking the spacebar key. The repetition priming test consisted of 240 masked trials presented subliminally, immediately followed by a word in lowercase letters, which was presented supraliminally (the ‘target’). Participants were asked to judge, in each trial, whether the target word was “natural” or “artificial”, using the left and right arrow keys on the keyboard, respectively. The target remained on the screen until one of these buttons was pressed. Critically, in half of the trials the masked word (the ‘prime’), presented in uppercase letters, was the same as the target at the end of the sequence (e.g., “MIRROR” followed by “mirror”). Therefore, if the masked word was indeed processed, response for the target should be facilitated, even though the two words were not presented in the same case, indicating semantic priming ^31^.

Response category (i.e., “natural” or “artificial”) was counterbalanced across targets within repeating and within non-repeating trials. Trials were ordered so that the response category did not repeat for more than three consecutive trials, and neither primes nor targets repeated in consecutive trials. As in Dehaene et al. (2001), the words in non-repeating trials always came from different response categories (e.g., masking “*WATER*”, which is natural, and then presenting “*broom*”, which is artificial). Lastly, words in non-repeating trials were of the same number of letters and did not share a first letter.

#### 8) Individualized Calibration Validation

We repeated a variation of the Individualized Calibration phase again at the end of the study. Trial and task structure were the same as in the original Individualized Calibration phase. This phase consisted of 40 trials: five trials using the blank duration used for the C-cued condition: 160 ms, 15 trials using the identified perceptual threshold (i.e., the blank duration used for the U-cued condition), 10 trials with a blank duration that was 13.33 ms longer than the perceptual threshold, and 10 trials with a blank duration that was 13.33 ms shorter than the perceptual threshold. Trial order was randomized. Two participants failed to complete the Individualized Calibration Validation phase due to their experiment taking more time than scheduled, and their data from this phase was not included in the analysis.

### Analyses

All analyses were completed using Matlab 2020a (MathWorks Inc., Natick, MA). Statistical analyses were conducted using linear mixed-effects models (*fitglme* function in Matlab), including a random intercept for participants. *F*-test and *t*-tests were used to analyze fixed effects (using the *anova* function of the *GeneralizedLinearMixedModel* class in Matlab).

Accuracy in Individualized Calibration and Individualized Calibration Validation trials was registered by the experimenter online after each trial. In five cases (0.1% of trials), the experimenter indicated that they were not sure about the participant’s last response, making the program ignore the last trial.

To mitigate spelling challenges, only the first 4 letters of each word were required in Recall. Responses were evaluated manually offline. Nonetheless, there were still twelve cases that were considered spelling mistakes (e.g., ‘jeas’ instead of ‘jeans’ and ‘herm’ instead of ‘harm’ for ‘harmonica’) and one case of a naming mistake (‘plan’ presumably for ‘plane’, instead of ‘airplane’) that were considered a correct response.

### Trial and Participant Exclusion

This study sought to investigate the effect of unconscious reactivation on pre-existing memories. Memories that were not properly formed at the onset, as indicated by poor memory immediately after learning and prior to reactivation, were thus excluded from analysis. Triads containing objects that were positioned over 250 pixels away from their veridical position during the Pre-reactivation Memory Test (2.7 inch; ∼35% of circle diameter and 2.5 times the allowed margin of error during Training) were removed from analysis. This led to the exclusion of M = 5.83, SD = 6.76 triads per participant (10.8%).

In addition to triad exclusions, three participants (6.25%) were excluded due to pre-reactivation memory score indicating lack of initial learning. To detect these participants, a two-parameter Weibull distribution was fit to each participant’s pre-reactivation error scores (in pixels). The Weibull describes a non-negative distribution of values that may be skewed. Pre-reactivation spatial memory accuracy should rise sharply on the left and have a long rightward tail, indicating that the majority of items are remembered well (i.e., with little spatial error) while a minority of items are weaker. We compared the fitted shape parameter of the distribution to the group average. The shape value of three participants (M = 288.62 SD = 54.23) was larger by more than two standard deviations from the group average (M = 115.47 SD = 55.04), indicating that the distribution of spatial error of these participants was abnormally centered around higher values, with tails on both sides of the center. Indeed, the percentage of outlier items of these participants (M = 51.23% SD = 15.42) was more than two standard deviations larger than the group average (M = 10.8% SD = 12.53). Thus, these participants were excluded from analysis due to overall poor memory performance.

In the Repetition Priming Test, incorrect responses (5.7% of responses), as well as responses that were either longer or shorter than each participant’s average by more than 2.5 standard deviations (2.6% of responses), were excluded from analysis. Additionally, participants whose responses were either longer or shorter than the group average by more than 2.5 standard deviations were removed from analysis (2/41 participants).

## Results

### Individualized visibility calibration

Participants first completed a task aimed at identifying their individual perceptual thresholds for being able to read masked words. This calibration converged within 81.6 trials on average (SD = 25.4; range: 53-145 trials). The average detected threshold was 33.82 ms (SD = 8.99 ms). Average accuracy in naming the masked words when exposed at the perceptual threshold was 3.1% (SD = 3.4%; Figure 2A). The average accuracy when using the maximal duration of 160 ms was 97.6% (SD = 10.9%).

When we checked perceptual thresholds again at the end of the session, average accuracy was higher than it was originally (M = 8.9%, SD = 10.9%; *t*(38) = −3.35, *p* = 0.002; Figure 2A). Importantly, however, performance remained far below the level of C-cued and under the 10% criterion (used to determine the threshold). Therefore, despite improvement in masked word identification, the U-cued presentation procedure did not reliably produce conscious perception either at the beginning or end of the session.

### Subliminal presentation and unconscious processing

To validate that our method of subliminal presentation produced unconscious processing, a Repetition Priming task was included. On each trial, participants made a judgement (natural/artificial) on a lower-case word preceded by a masked upper-case word (same parameters as in main tasks), which, on half the trials, was the same word. Results revealed a significant priming effect, in that responses to target words were significantly faster when the preceding masked word was the same compared to different [*t*(38) = 2.23, *p* = 0.032; Figure 2B]. This manipulation check replicated previously reported unconscious repetition priming (e.g., ^31^ and demonstrated that the subliminal presentation procedure was potent enough to affect semantic processing.

### Training

The main memory task used in this study involved triadic associations among adjectives, objects, and on-screen positions. Memories were reactivated for items in six of the nine categories: three were unconsciously reactivated and three were consciously reactivated. Only half of the triads within each of these categories were cued (U-cued; C-cued), whereas the others were not (U-noncued; C-noncued). In the remaining three categories, none of the triads were reactivated (NR). Triads were assigned to conditions according to performance during training, such that memory strength would be balanced across conditions (see *Procedure*). Indeed, all pre-manipulation performance measures, including the number of learning iterations required to reach learning criterion, the average success rate of adjective-object association during training, the average error in object-position association during training, and the object-position error in Pre-reactivation Memory Test, were statistically equivalent across conditions (Table 1).

**Table 1.** Pre-manipulation performance measures of associative triads, per condition (mean ± SD).

|  | NR | C-cued | C-noncued | U-cued | U-noncued | Condition effect |
| --- | --- | --- | --- | --- | --- | --- |
| Number of required training iterations | 3.7 ± 1.1 | 3.8 ± 1.1 | 3.7 ± 1.2 | 3.6 ± 1.1 | 3.7 ± 1.4 | $F(4, 2042) = 0.39, p = 0.816$ |
| Training object naming accuracy | 90.4 ± 6.0 | 89.9 ± 7.2 | 91.2 ± 6.3 | 90.4 ± 6.8 | 91.8 ± 5.5 | $F(4, 2042) = 1.25, p = 0.289$ |
| Training average position error | 71.5 ± 21.9 | 72.5 ± 24.7 | 77.7 ± 25.7 | 70.8 ± 23.6 | 75.8 ± 26.3 | $F(4, 2042) = 1.58, p = 0.176$ |
| Pre-reactivation object naming accuracy | 82.3 ± 16.0 | 84.0 ± 19.3 | 84.0 ± 17.9 | 85.4 ± 15.7 | 87.3 ± 16.5 | $F(4, 2042) = 1.38, p = 0.239$ |
| Pre-reactivation position error | 69.7 ± 15.3 | 71.7 ± 16.2 | 74.8 ± 18.0 | 71.9 ± 20.1 | 73.4 ± 20.4 | $F(4, 2042) = 0.85, p = 0.496$ |

### Accessibility of proximal associations

We first assessed the effect that reactivation of adjectives (e.g., “SCARED”) had on recall of objects linked with that adjective (e.g., “banana”). Participants were asked to recall as many objects as they could for each semantic category (e.g., “apple”, “banana”, “grapes”, etc.). Objects were divided into five conditions: unconsciously cued (U-cued); noncued members of an unconsciously cued category (U-noncued); consciously cued (C-cued); noncued members of a consciously cued category (C-noncued); non-reactivated (NR; see Figure 1B). A main effect of condition was found in the probability to recall objects (*F*(4, 1994) = 4.03, *p* = 0.003). Follow-up analysis revealed that recall of C-cued objects was significantly higher than recall of objects of all other conditions (C-cued vs. NR: *t*(1994) = −2.75, *p* = 0.006; C-cued vs. U-noncued: *t*(1994) = - 3.32, *p* = 0.001; C-cued vs. U-cued: *t*(1994) = −3.18, *p* = 0.001; C-cued vs. C-noncued: *t*(1994) = - 3.32, *p* = 0.001; Figure 2C). This indicates that consciously reactivating a triad memory by presented the adjective cue led to greater accessibility of the associated object. However, the reactivation of some triads from a semantic group had no effect on other triads from the same group, as recall rates of C-noncued triads was similar to that of other noncued items (C-noncued vs. NR: *t*(1994) = 1.08, *p* = 0.28; C-noncued vs. U-noncued: *t*(1994) = 0.02, *p* = 0.984). Finally, recall rates of U-cued items were similar to those of noncued items (U-cued vs. NR: *t* (1994) = 0.91, *p* = 0.362; U-cued vs. U-noncued: *t*(1994) = −0.13, *p* = 0.896; U-cued vs. C-noncued: *t*(1994) = −0.15, *p* = 0.881). These results additionally serve to validates our manipulation by demonstrating that triads in the U-cued condition were not consciously perceived, since their effects differ from those in the C-cued condition.

### Accessibility of distal associations

Subsequent analyses focused on memory for more ‘distal’ parts of the triadic memory, namely, the effects of reactivation on memory for the associated object positions. No main effect of condition was found in post-reactivation positioning error (*F*(4, 2042) = 1.61, *p* = 0.168), but planned comparisons revealed that post-reactivation positioning error of C-noncued items was higher than that of U-noncued items (*t*(40) = 1.90, *p* = 0.033, one-tailed) and marginally higher than NR items (*t*(40) = 1.55, *p* = 0.064, one-tailed), supporting a consciousness-driven inhibition effect on these spatial associations (Figure 2D).

As previous studies have suggested that the benefits of reactivation during sleep and awake states differs based on initial memory strength, with preferential benefits to weak ^12,25,26^ and intermediate ^32,33^ strengths, we next incorporated initial memory strength in our model of post-reactivation spatial error. Despite being trained to reach the same learning criterion, triads in our task varied in how strongly they were encoded, as reflected in the distribution of pre-reactivation spatial error values (M = 71.8 pixels, SD = 49; Figure 3A). Thus, we considered pre-reactivation spatial error as the metric of memory strength in our task. This model revealed a main effect of pre-reactivation error (*F*(1, 2037) = 227.34, *p* < 0.001), a main effect of condition (*F*(4, 2037) = 3.08, *p* = 0.015), and an interaction between them (*F*(4, 2037) = 2.99, *p* = 0.018; Figure 3A).

**Figure 3.**
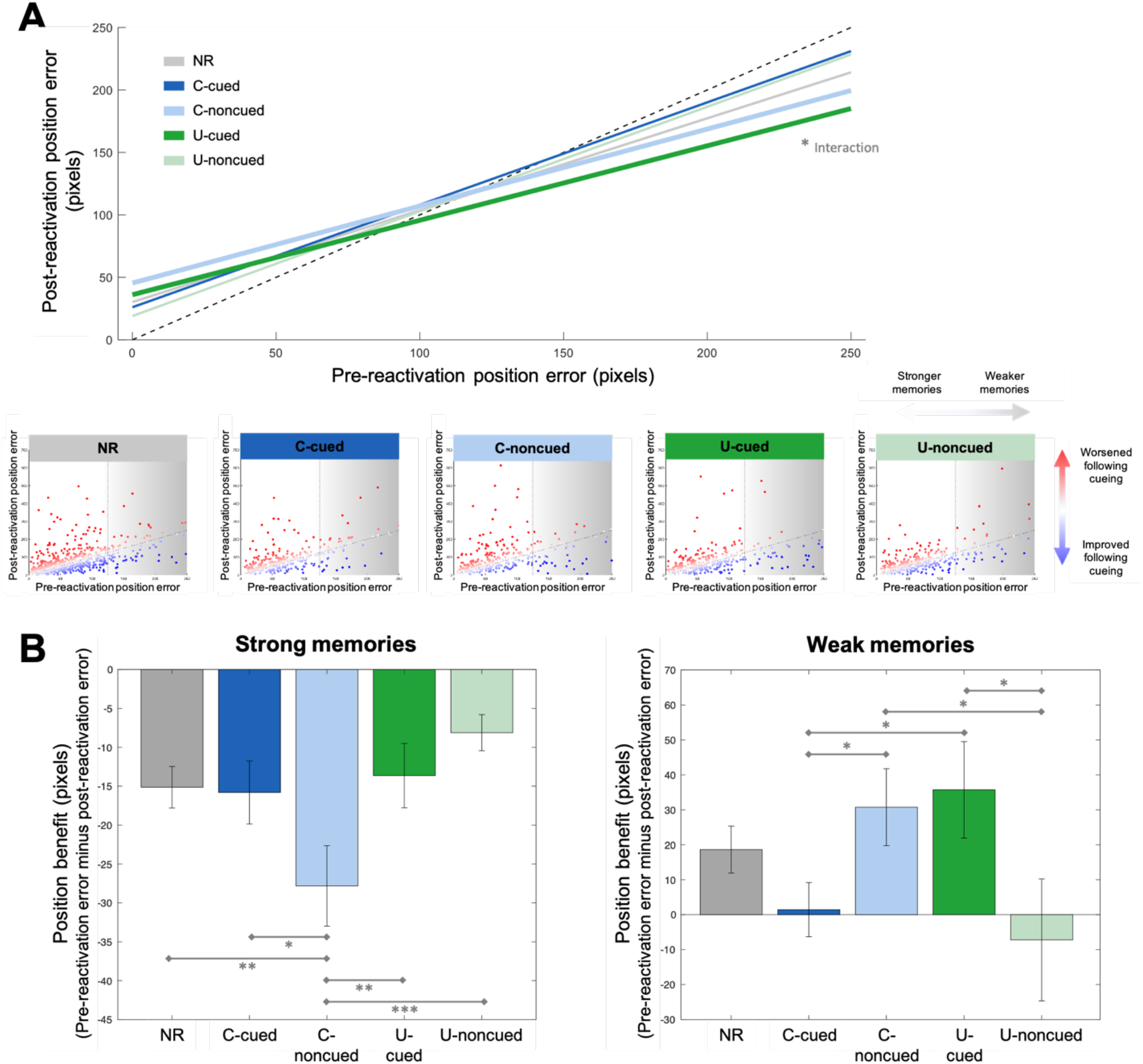
Spatial memory after reactivation. **A) Top panel:** Stronger memories (with small initial positioning error) remained stronger, and weaker memories remained weaker over time. However, weak U-cued and C-noncued memories benefited from reactivation more than other weak memories. **Bottom panel:** Scatter plots of individual triads depict the relationship between pre- and post-reactivation spatial memory for all objects in each condition. Weak memories from the U-cued and C-noncued conditions mostly improved from pre- to post-reactivation. Dashed lines represent the *y* = *x* line in all panels. **B) Left panel:** Memory for the position of objects from strongly learned memories deteriorated over time. Memory for C-noncued triads deteriorated more than memory for triads of all other conditions. **Right panel:** Weak memories benefited from reactivation in a condition-dependent manner. U-cued and C-noncued memories benefited more than memories of some of the other conditions. Error bars reflect SEM. NR – non-reactivated; C-cued – consciously cued; C-noncued – noncued member of a list with C-cued items. U–cued unconsciously cued; U-noncued – noncued member of a list with U-cued items. * *p* < 0.05, ** *p* < 0.01, *** *p* < 0.001.

A main effect of pre-reactivation error can be expected, as weak memories remain relatively weaker by the end of the experiment, and strong memories remain relatively stronger. Followup analyses on the main effect of condition demonstrated that, when taking pre-reactivation position error into account, post-reactivation position error of C-noncued items was significantly higher than that of NR, U-noncued and C-cued items (C-noncued vs. NR: *t*(2037) = - 2.19, *p* = 0.029; C-noncued vs. U-noncued: *t*(2037) = −3.26, *p* = 0.001; C-noncued vs. C-cued: *t*(2037) = −2.40, *p* = 0.017). Further, U-cued items had higher post-reactivation position error than U-noncued items (*t*(2037) = −2.10, *p* = 0.036). Follow-up analyses of the interaction effect revealed that memory-strength-dependent effects of U-cued spatial memories differed from all other conditions except C-noncued (U-cued vs. NR: *t*(2037) = 1.75, *p* = 0.079; U-cued vs. U-noncued: *t*(2037) = 2.62, *p* = 0.009; U-cued vs. C-cued: *t*(2037) = 2.48, *p* = 0.013; U-cued vs. C-noncued: *t*(2037) = 0.22, *p* = 0.823). These effects are driven by improvements for weaker memories in the U-cued group (and detriments for stronger ones) relative to the other conditions. A similar effect was found for C-noncued memories (C-noncued vs. C-cued: *t*(2037) = 2.21, *p* = 0.027; C-noncued vs. U-noncued: *t*(2037) = 2.37, *p* = 0.018), although here no difference was found with non-reactivated categories (*t*(2037) = 1.47, *p* = 0.141). There was no difference between C-noncued and U-cued items in the main effect of memory strength (*t*(2037) = −1.17, *p* = 0.243), nor was there a difference in their interactions with memory strength (*t*(2037) = −0.22, *p* = 0.823). Therefore, the weaker a spatial memory was initially (i.e., with larger pre-reactivation errors), the more it improved, and this effect was similar in U-cued and C-noncued conditions and stronger than in other conditions.

Having established an encoding-strength difference in the effect of reactivation type, we next analyzed the benefit that reactivation had on memories for strong and weak memories separately (Figure 3B). The pre-reactivation error range was split in two, so that triads were considered “strong” if pre-reactivation error was under or equal to 125 pixels (half the allowed error, see *Trial and Participant Exclusion*) and “weak” if it was above 125 pixels (Figure 3B). On average, 13.6% ± 7.5% of a participants’ triads were considered weak according to this division. A position benefit score was defined per triad as the change in position error from pre- to post-reactivation [i.e., (Pre-Position ↔ True-Position) – (Post-Position ↔ True-Position)]. A positive value implies that spatial memory had improved following reactivation, and a negative value implies that the memory had degraded following reactivation.

For strong memories, the position error increased in all conditions, meaning that forgetting was dominant and not influenced by reactivation (Figure 3B, left panel). This is to be expected and reflects a trivial regression to the mean. However, a main effect of condition (*F*(4, 1771) = 4.37, *p* = 0.002) indicated that memory decline was not equivalent across all conditions. C-noncued triads grew worse in spatial memory precision as compared to all other conditions (C-noncued vs. NR: *t*(1771) = 2.87, *p* = 0.004; C-noncued vs. C-cued: *t*(1771) = 2.29, *p* = 0.022; C-noncued vs. U-cued: *t*(1771) = 3.05, *p* = 0.002; C-noncued vs. U-noncued: *t*(1771) = 4, *p* < 0.001) even though pre-reactivation positioning error of strong memories was equivalent among all conditions (*F*(4, 1771) = 0.48, *p* = 0.753). These results suggest that for strong memories, reactivation had a detrimental effect on spatial memory for related memories when reactivation was conscious, but not when it was unconscious. This finding resonates with the phenomenon of retrieval-induced forgetting ^14^, in which reactivation of a memory leads to inhibition of related memories ^20^. The pattern of results found in position memory is aligned with our hypothesis that conscious reactivation of a target item is accompanied by inhibition, while unconscious reactivation does not. However, it should be noted that there were no apparent benefits for cueing either consciously or unconsciously (C-cued vs NR: *t*(1771) = 0.19, *p* = 0.848; U-cued vs NR: *t*(1771) = −0.65, *p* = 0.514), nor were there differences between the benefits for C-cued relative to U-cued items (C-cued vs U-cued: *t*(1771) = 0.73, *p* = 0.468).

Within the weak memories, benefits of reactivation were evident (Figure 3B, right panel). Once again, these benefits, or average, are to be expected based on the selection of weak memories and given the expected regression to the mean. Critically, however, results show a main effect of condition for these memories (*F*(4, 266) = 2.47, *p* = 0.045), even though pre-reactivation errors were the same in weak memories across conditions (*F*(4, 266) = 1.16, *p* = 0.327). Weak U-cued items improved more than weak C-cued items (*t*(266) = −1.99, *p* = 0.048) and weak U-noncued (*t*(266) = −2.22, *p* = 0.027). A similar pattern of results was also found for weak C-noncued items (C-noncued vs. C-cued: *t*(266) = −2.20, *p* = 0.029; C-noncued vs. U-noncued: *t*(266) = −2.43, *p* = 0.016). Reactivation benefits for weak U-cued and weak C-noncued, however, were equivalent (*t*(266) = 0.21, *p* = 0.837). These results suggest, again, that subliminal activation, either via unconscious reactivation (i.e., U-cued) or via spreading activation from conscious reactivation(i.e., C-noncued), preferentially benefited weak memories.

### Reorganization of distal associations

Finally, we investigated whether systematic biases existed in post-reactivation spatial memory. An implication of our hypothesis that unconscious reactivation involves lesser inhibition is that spreading activation will not be countered by inhibition, and so unconsciously reactivated memories will co-activate with related memories. Such co-activation may result in integration in memory ^29^. Thus, in our experiment, U-cued items may be active alongside U-noncued items, and the two could be bound together, more than C-cued and C-noncued items would. To test this, the spatial layout of items on the grid was set so that cued and noncued items of each category occupied separate neighboring quadrants (Figure 1B). If indeed U-noncued items were activated alongside U-cued items, we would expect the reported position memory of U-noncued items to be biased towards the positions of U-cued items.

To examine this hypothesis, we calculated the mean position of cued and noncued members of each category, and termed these the “centers” of the cued and the noncued quadrants. We then compared the distance between where objects were placed and the quadrant centers pre-reactivation and post-reactivation [i.e., (Post-Position ↔ Quadrant center) – (Pre-Position ↔ Quadrant center)]. The difference between the two is a measure of shift towards the quadrant center (negative values indicating a shift towards the center and positive values a shift away from it). Our analysis considered two types of shifts: noncued items that shifted toward the cued center (i.e., U-noncued → U-cued center; C-noncued → C-cued center), and cued items that shifted toward the noncued center (i.e., U-cued → U-noncued center; C-cued → C-noncued center).

For noncued items, we found an interaction between condition and strength in accounting for the shift towards the cued quadrant (*F*(1, 681) = 8.70, *p* = 0.003; There was no main effect of condition: *F*(1, 681) = 0.01, *p* = 0.937, nor a main effect of strength: *F*(1, 681) = 1.60, *p* = 0.207). Follow-up analysis of weak memories only revealed that, post-reactivation, weak U-noncued items shifted closer to the quadrant of their U-cued peers more than weak C-noncued items shifted towards the quadrant of their C-cued peers (*t*(89) = 2, *p* = 0.048; Figure 4). This effect was not present for strong memories (*t*(592) = 0.09, *p* = 0.927). We conducted a permutation test to validate this finding. The same analysis was applied to 10,000 random shufflings of labels among conscious and unconscious category labels (i.e., randomly selecting three categories as conscious and three as unconscious among the six categories that were not NR per each participant). The coefficient size obtained in our data (*β* = 38.61) was more extreme than 96.4% of the coefficients obtained under random permutations (M = 0.19, SD = 18.72; *p* = 0.036), suggesting that the unconscious nature of cueing had indeed introduced a systematic bias towards the cued positions. In cued items, no difference was found between the conditions in their shift towards their noncued quadrants (main effect of condition: *F*(1, 683) = 1.02, *p* = 0.313; main effect of strength: *F*(1, 683) = 0.19, *p* = 0.660; interaction between condition and strength: *F*(1, 683) = 0.09, *p* = 0.766).

**Figure 4.**
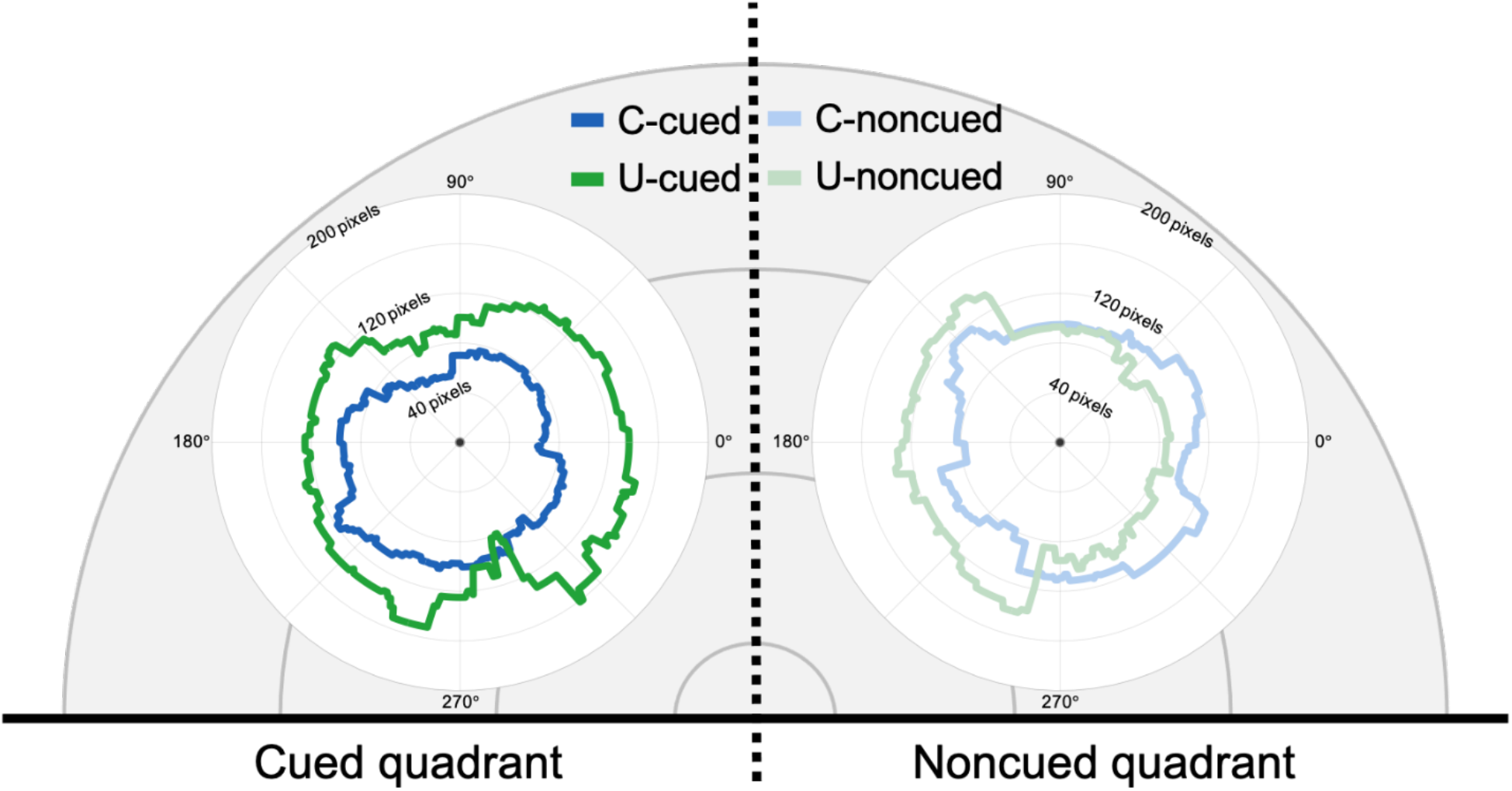
Shifts in spatial error of weak memories post-reactivation. For this visualization, item positions were transformed so that categories were superimposed over the top 180° of the grid, with the cued quadrant on the left and the non-cued quadrant on the right. Lines indicate the direction (in degrees) and size (in pixels) of the difference between post-reactivation and pre-reactivation positioning (smoothed using a sliding average over 145°). This visualization shows a leftwards shift in weak U-noncued positioning post-reactivation, i.e., that these items were systematically placed closer to the cued quadrant of the same category after reactivation.

## Discussion

In this paper, we contrasted the effects of awake conscious and unconscious reactivation on memory. Participants were trained to learn triadic memories (adjective-object-position) prior to reactivation when a portion of these memories were then cued by presenting the adjective either subliminally or supraliminally. We hypothesized that conscious and unconscious memory reactivation would benefit memory in distinct ways owing to the observation that, unlike unconscious reactivation, conscious reactivation should be accompanied by more inhibition of other related memories ^14,21^. Therefore, we predicted that conscious reactivation would strongly benefit the consciously retrieved or rehearsed content but may also have detrimental effects on semantically related non-reactivated memories, as observed in RIF experiments ^14^. By contrast, we predicted that unconscious reactivation would be more likely to improve memory for weaker associations that are more distant to the cued adjective, in this case the spatial position of the cued triad, without causing impairment to associated memories. We also tested an additional implication of our hypothesis, that unconscious reactivation would promote integration of associated memories due to nonrestrictive spreading activation.

Results provided partial support for our hypotheses. As predicted, conscious reactivation improved retrieval of the cued objects compared to unconscious reactivation. With regards to the more distal spatial position association, conscious and unconscious cueing had a memory-strength-dependent effect that differed between conditions. Consciously reactivating some memories from a semantic category affected other memories belonging to that category according to their strength: strongly encoded memories were impaired while weakly encoded memories were improved (more than can be expected by regression to the mean). Unconscious reactivation, on the other hand, benefited weakly encoded memories and caused their integration with associated memories, without impacting strongly encoded memories. Thus, these results highlight that conscious and unconscious memory reactivation have qualitatively distinct consequences on memory strength and integration.

The detrimental effect that conscious but not unconscious reactivation had on related memories that were strongly encoded is in line with our hypothesis of decreased inhibition recruited by unconscious reactivation. The RIF effect is mostly driven by inhibitory dynamics ^20^ and is thus expected to be found following conscious reactivation but not following unconscious reactivation. The finding that only strongly encoded memories suffered impairment following conscious cueing is consistent with the competitive framework of conscious retrieval and the ‘interference dependence’ aspect of RIF, which demonstrates that related memories are suppressed according to the strength of their link to the reactivated memory ^14^. Weakly encoded memories may also be weakly linked with the reactivated category. They could thereby be rescued from RIF, perhaps due to nonmonotonic plasticity dynamics ^34^.

Beneficial effects of reactivation on weakly encoded memories are in line with previous reports on the contribution of offline reactivation to memory. For example, Schapiro et al. (2018) showed that memory reactivation during post-task rest predicted memory improvement, but only for weakly encoded memory. Similarly, targeted memory reactivation studies have shown a selective benefit for weakly encoded memories during sleep ^25–28^ and during wakefulness ^12^. Importantly, these effects cannot be explained by regression to the mean or floor effects, since they contrast weakly encoded memories that were either reactivated or not and shared similar starting points and expected trajectories. This selective benefit is consistent with neurobiologoical models of synaptic tag and capture ^35,36^ and may be the result of prioritized offline consolidation of memories that have been tagged as needing such strengthening ^37,38^. The effects of unconscious reactivation on weakly, but not strongly, encoded memories are therefore in line with emerging work examining the effects of exogenous or endogenous reactivation on later memory.

Lastly, unconscious reactivation did not affect the accessibility of other semantically related memories but it did affect their content, indicating some spreading activation in unconscious reactivation, as is also found during sleep ^28^. Furthermore, spatial memory of cued items seemed to infiltrate spatial memory of their noncued associates, supporting the hypothesis that unconscious reactivation would permit their co-activation ^29^. This integration resonates with the emergance of relational memory found following sleep consolidation ^39^ and increases in neural integration ^40^. However, more work is needed to understand this aspect of unconscious reactivation, and whether it has similar influences on memory during sleep and wakefulness. Unconscious reactivation during wake and sleep may have distinct influences on memory, and unconscious activations during sleep may also be stronger than the activations used in this task, which were set to be weak in order to remain unconscious.

Results did not meet our hypotheses in three ways. First, conscious reactivation had no determinantal effect on the recall of objects from the same category that were not cued, as would be predicted by RIF. This lack of an RIF effect in our first-order association may be due to category binding being weaker in our design than in typical RIF studies, in which the category word is always presented alongside the cue word during both study and retrieval. This could have caused the categorical effect to be too weak to affect object recall, and so only discernable in the more sensitive measure of spatial memory. Second, conscious reactivation had no effect on spatial precision of the target memory, but rather only on its associated memories from the same category. Third, the benefits of unconscious reactivation for weakly encoded memories were not significantly larger than the effects observed in non-reactivated memories. These last two null-effects are hard to interpret. It may be that even though differences between unconsciously-reactivated and non-reactivated items were clear in regression analyses, our design was underpowered to detect these effects in aggregated spatial error. Future studies should be powered based on the effects observed in this study to shed light on these findings.

The key feature of the current work was our intentional manipulation of consciousness. As discussed above, several studies have considered the role of offline memory processing during wakefulness ^3,8–11,41^; see Wamsley (2022) for review). However, although offline states were linked before to sensory decoupling and lack of attention to reactivated material ^5^, no study to the best of our knowledge has examined the effect that conscious awareness of the reactivated material has on subsequent memory. Tambini and colleagues (2017) set the stage for isolating the effects of *unaware a*wake processing on memory. Our experiment has taken this question further by examining *unconscious a*wake processing, guided by a novel hypothesis that processing outside of conscious access has unique characteristics that may be beneficial for certain mnemonic operations. While the impact of consciousness on memory accessibility has been extensively studied (see, e.g., ^42^), our study investigates the effect that unconscious reactivation has on later memory accessibility and integration at a later timepoint. Hence, it could not be attributed to mere short-term facilitation, as effects of unconscious priming are typically short-lived, on the order of a few hundred milliseconds ^43,44^. Rather, they suggest longer-term modification in the storage of reactivated memories ^45^.

Nevertheless, our design had some notable limitations. First, our reactivation paradigm relied on links between the adjectives (that served as cues), the objects, and finally their spatial positions (which was the measured memory). Adjective-object associations were well-learned before reactivation (∼84% correctly recalled), while the second-order object-position associations were purposefully not learned to ceiling in order to allow for variability in initial memory strength because the extant literature strongly suggest that offline reactivation may selectively benefit weaker memories. However, one consequence of this experimental choice is that the initial memory links may not have been sufficiently strong to reliably reactivate downstream spatial positions given the adjective cue. Second, our unconscious reactivation trials differed from those used by Tambini and colleagues (2017) in a way that may have impacted their effects. Whereas the previous study presented cues while participants were engaged in a boring, mind-wandering-inducing task, our design mixed conscious and unconscious reactivation trials within the same blocks, thereby requiring participants to be constantly alert and attending to the presented information. This may have reduced the effectiveness of the unconscious manipulation, that may depend on the participants being in an ideal ‘offline’ state during reactivation ^12,41^.

By contrasting conscious and unconscious reactivation within the same design, our results constitute the first step in considering the role of consciousness in associative spread and the ensuing results on memory accessibility and organization. Indeed, our results indicate that consciousness plays an important role in determining these trajectories. While conscious rehearsal remains more advantageous for retrieval later in time, our study demonstrates that unconscious reactivation also has consequences for memory representations, which are different, and probably more similar to the effects of endogenous memory replay during sleep or rest. To fully understand the impact of different consciousness states on consolidation, future studies should examine unconscious reactivation induced by different means (e.g., attentional blink, continuous flash suppression), as they have been shown to be processed differently ^46^, as well as the full range of naturally occurring consciousness states, including full alertness, mind wandering, and the different stages of sleep.

## Acknowledgements

This work was supported by National Institute of Health (USA) grant R00-MH122663 for ES, National Science Foundation (USA) grant BCS-2048681 for KAP, and NSF-BSF Grant 2048587

## References

1. Paller, K. A. & Wagner, A. D. Observing the transformation of experience into memory. Trends Cogn Sci 6, 93–102 (2002).

2. Squire, L. R., Cohen, N. J. & Nadel, L. The medial temporal region and memory consolidation: a new hypothesis. in Memory Consolidation: Psychobiology of Cognition (eds. Weingartner, H. & Parder, E. S.) 185–210 (1984).

3. Tambini, A. & Davachi, L. Awake Reactivation of Prior Experiences Consolidates Memories and Biases Cognition. Trends Cogn Sci 23, 876–890 (2019).

4. Diekelmann, S. & Born, J. The memory function of sleep. Nat Rev Neurosci 11, 114–126 (2010).

5. Wamsley, E. J. Offline memory consolidation during waking rest. Nature Reviews Psychology 0123456789, (2022).

6. Staresina, B. P. et al. Hierarchical nesting of slow oscillations, spindles and ripples in the human hippocampus during sleep. Nature Neuroscience 2015 18:11 18, 1679–1686 (2015).

7. Staresina, B. P., Niediek, J., Borger, V., Surges, R. & Mormann, F. How coupled slow oscillations, spindles and ripples coordinate neuronal processing and communication during human sleep. Nature Neuroscience 2023 1–9 (2023).

8. Tambini, A. & Davachi, L. Persistence of hippocampal multivoxel patterns into postencoding rest is related to memory. Proc Natl Acad Sci U S A 110, 19591–19596 (2013).

9. Tambini, A., Ketz, N. & Davachi, L. Enhanced Brain Correlations during Rest Are Related to Memory for Recent Experiences. Neuron 65, 280–290 (2010).

10. Schapiro, A. C., McDevitt, E. A., Rogers, T. T., Mednick, S. C. & Norman, K. A. Human hippocampal replay during rest prioritizes weakly learned information and predicts memory performance. Nature Communications 2018 9:1 9, 1–11 (2018).

11. Staresina, B. P., Alink, A., Kriegeskorte, N. & Henson, R. N. Awake reactivation predicts memory in humans. Proc Natl Acad Sci U S A 110, 21159–21164 (2013).

12. Tambini, A., Berners-Lee, A. & Davachi, L. Brief targeted memory reactivation during the awake state enhances memory stability and benefits the weakest memories. Sci Rep 7, 1–17 (2017).

13. Healey, M. K., Campbell, K. L., Hasher, L. & Ossher, L. Direct Evidence for the Role of Inhibition in Resolving Interference in Memory. Psychol Sci 21, 1363–1547 (2010).

14. Anderson, M. C. Rethinking interference theory: Executive control and the mechanisms of forgetting. J Mem Lang 49, 415–445 (2003).

15. Kuhl, B. A., Bainbridge, W. A. & Chun, M. M. Neural reactivation reveals mechanisms for updating memory. Journal of Neuroscience 32, 3453–3461 (2012).

16. Tal, A. & Bar, M. The proactive brain and the fate of dead hypotheses. Front Comput Neurosci 8, 1–6 (2014).

17. Roediger, H. L. & Karpicke, J. D. Test-enhanced learning: Taking memory tests improves long-term retention. Psychol Sci 17, 249–255 (2006).

18. Hulbert, J. C. & Norman, K. A. Neural differentiation tracks improved recall of competing memories following interleaved study and retrieval practice. Cerebral Cortex 25, 3994– 4008 (2015).

19. Kuhl, B. A., Dudukovic, N. M., Kahn, I. & Wagner, A. D. Decreased demands on cognitive control reveal the neural processing benefits of forgetting. Nat Neurosci 10, 908–914 (2007).

20. Storm, B. C. et al. A review of retrieval-induced forgetting in the contexts of learning, eyewitness memory, social cognition, autobiographical memory, and creative cognition. Psychology of Learning and Motivation - Advances in Research and Theory 62, 141–194 (2015).

21. Schmitz, T. W., Correia, M. M., Ferreira, C. S., Prescot, A. P. & Anderson, M. C. Hippocampal GABA enables inhibitory control over unwanted thoughts. Nat Commun 8, 1–11 (2017).

22. Hu, X., Cheng, L. Y., Chiu, M. H. & Paller, K. A. Promoting memory consolidation during sleep: A meta-analysis of targeted memory reactivation. Psychol Bull 146, 218–244 (2020).

23. Oudiette, D. & Paller, K. A. Upgrading the sleeping brain with targeted memory reactivation. Trends Cogn Sci 17, 142–149 (2013).

24. Alm, K. H., Ngo, C. T. & Olson, I. R. Hippocampal signatures of awake targeted memory reactivation. Brain Struct Funct 224, 713–726 (2019).

25. Cairney, S. A., Lindsay, S., Sobczak, J. M., Paller, K. A. & Gaskell, M. G. The benefits of targeted memory reactivation for consolidation in sleep are contingent on memory accuracy and direct cue-memory associations. Sleep 39, 1139–1150 (2016).

26. Creery, J. D., Oudiette, D., Antony, J. W. & Paller, K. A. Targeted memory reactivation during sleep depends on prior learning. Sleep 38, (2015).

27. Schechtman, E. et al. Sleep reactivation did not boost suppression-induced forgetting. Sci Rep 11, (2021).

28. Schechtman, E., Heilberg, J. & Paller, K. A. Memory consolidation during sleep involves context reinstatement in humans. CellReports 42, (2023).

29. Lewis, P. A. & Durrant, S. J. Overlapping memory replay during sleep builds cognitive schemata. Trends Cogn Sci 15, 343–351 (2011).

30. Biderman, D., Shir, Y. & Mudrik, L. B or 13? Unconscious Top-Down Contextual Effects at the Categorical but Not the Lexical Level. Psychol Sci 31, 663–677 (2020).

31. Dehaene, S. et al. Cerebral mechanisms of word masking and unconscious repetition priming. Nat Neurosci 4, 752–8 (2001).

32. Schechtman, E., Heilberg, J. & Paller, K. A. Context matters: changes in memory over a period of sleep are driven by encoding context. Learning & Memory 30, 36–42 (2023).

33. Stickgold, R. How do I remember? Let me count the ways. Sleep Med Rev 13, (2009).

34. Ritvo, V. J. H., Turk-Browne, N. B. & Norman, K. A. Nonmonotonic Plasticity: How Memory Retrieval Drives Learning. Trends Cogn Sci 23, 726–742 (2019).

35. Redondo, R. L. & Morris, R. G. M. Making memories last: the synaptic tagging and capture hypothesis. Nature Reviews Neuroscience 2011 12:1 12, 17–30 (2010).

36. Frey, U. & Morris, R. G. M. Synaptic tagging and long-term potentiation. Nature 1997 385:6616 385, 533–536 (1997).

37. Dunsmoor, J. E., Murty, V. P., Clewett, D., Phelps, E. A. & Davachi, L. Tag and capture: how salient experiences target and rescue nearby events in memory. Trends Cogn Sci 26, 1–14 (2022).

38. Lewis, P. A. & Bendor, D. How Targeted Memory Reactivation Promotes the Selective Strengthening of Memories in Sleep. Current Biology 29, R906–R912 (2019).

39. Ellenbogen, J. M., Hu, P. T., Payne, J. D., Titone, D. & Walker, M. P. Human relational memory requires time and sleep. Proc Natl Acad Sci U S A 104, 7723–7728 (2007).

40. Tompary, A. & Davachi, L. Consolidation Promotes the Emergence of Representational Overlap in the Hippocampus and Medial Prefrontal Cortex. Neuron 96, 228-241.e5 (2017).

41. Nicosia, J. & Balota, D. A. Targeted memory reactivation and consolidation-like processes during mind-wandering in younger and older adults. J Exp Psychol Learn Mem Cogn (2022).

42. Churchill, L. Conscious and Unconscious Semantic Activation in Episodic Memory Retrieval. Washington University Undergraduate Research Digest 12, 12–20 (2017).

43. Kouider, S. & Dehaene, S. Levels of processing during non-conscious perception: A critical review of visual masking. Philosophical Transactions of the Royal Society B: Biological Sciences 362, 857–875 (2007).

44. Greenwald, A. G., Draine, S. C. & Abrams, R. L. Three Cognitive Markers of Unconscious Semantic Activation. Science (1979) 273, 1699–1702 (1996).

45. Reber, T. P. & Henke, K. Integrating unseen events over time. Conscious Cogn 21, 953– 960 (2012).

46. Breitmeyer, B. G. Psychophysical ‘blinding’ methods reveal a functional hierarchy of unconscious visual processing. Conscious Cogn 35, 234–250 (2015).

